# Cued replay of memory contents in human infants

**DOI:** 10.1101/2025.06.12.659246

**Authors:** Christopher M. Postzich, Johanna Finnemann, Michael A. Skeide

## Abstract

Sequentially structured and temporally compressed reactivation of memory contents plays a key role for learning in the mature mammalian brain. This type of neural coding, known as replay, could also drive the enormous progress in sequential learning seen in human early life. Rodent models, however, suggest that replay emerges slowly and refines gradually in childhood. To test whether this prediction holds in humans, we introduced a novel fast event-related stimulation paradigm for infants undergoing non-invasive electroencephalography. Leveraging time-resolved decoding techniques we discovered multimodal replay already in infants as young as 10 months. Reactivation responses revealed a forward sequential structure and temporal compression of the learnt sequence by a factor of 6 to 7. These results demonstrate cued replay activity at the macroscopic level in the first year of life, an age at which it has not been found in rodent place cells. Our insights stimulate future research exploring how replay contributes to the development of fundamental human cognitive capacities including language processing and action planning.

## Introduction

Electrophysiological recordings from hippocampal place cells revealed that rats reactivate sequences of previously visited locations during subsequent slow wave sleep and awake rest^1–3^. This so-called hippocampal replay follows two neural coding principles: First, it preserves the structure of the spatial sequence either in forward or backward order. Second, the duration of the spatial sequence is compressed in time by a factor of 5 to 20^1–3^. Hippocampal replay in rats has been associated with spatial memory consolidation and cognitive functions like spatial working memory and spatial action planning^4–6^. Multi-unit recordings have shown that replay is not limited to the hippocampal formation but occurs simultaneously in the visual cortex and that both regions reactivate the same episodes of past spatial experience^7^. In human adults, hippocampal and visual cortical replay has also been elicited by visual object stimuli embedded into non-spatial tasks, including sequential decision-making and rule learning^8–10^. A common approach in these experiments is to employ cues to trigger replay responses during awake rest that are large enough to be detectable with macroscale magnetencephalography and magnetic resonance imaging^11–13^.

Recent work in rats has suggested that replay-like neural coding slowly emerges only after postnatal day 20 (corresponding to late toddlerhood in humans) and still gradually refines at postnatal day 29 to 32 (corresponding to late childhood in humans)^14,15^. A similarly slow developmental trajectory might be expected in humans given that episodic memory formation in infants has been observed so far only in the context of basic recognition and association tasks^16–19^. At the same time, infants exhibit sequential learning in core domains of cognition, such as language processing or action planning^20–21^. It is plausible to hypothesize that such early forms of learning may be supported by replay. However, whether human replay is already operational in developing brains, let alone infant brains, is currently unknown.

To illuminate this question, we developed a fast-event related stimulus presentation design tailored to the time constraints of electroencephalographic recording in infants. In the first part of the experiment, the localizer part, participants viewed three different natural objects in random order, each of which was accompanied by a simultaneous low, middle or high tone. In the second part, the sequence learning part, the three visual objects and corresponding tones were presented in a fixed order to establish a sequential representation in memory. Finally, in the resting-state part, the low tone accompanying the first visual object in the previously established sequence was delivered to cue the reactivation of the whole sequence. The analytical key for detecting both sequential structure in and temporal compression was a combination of a one-vs-rest logistic regression classifier trained on the localizer data and a time-resolved decoding framework known as temporally delayed linear modeling. In this framework, sequences of stimulus probability transitions are modelled and tested while controlling for other possible sources of variance of the sequence and potential oscillatory confounds within the raw signal. This approach robustly revealed both the sequential structure and the temporal compression of replay across MEG studies in adults^9,12^.

Applying time-resolved decoding techniques to the electrophysiological time courses, recorded at a sampling rate of 1000 Hz with a low-pass filter of 250 Hz, we found time-compressed forward reactivation of the correctly-ordered sequence of previously seen objects in awake resting human infants. Cued replay compressed the duration of the original sequence by a factor of 6 to 7.

## Results

### Experimental stimulation design

To render it possible to detect replay in infants (see Supplementary Fig. S1a for participant characteristics) we designed an electroencephalography (EEG) experiment consisting of three stimulation parts. The first part serves as a functional localizer for object recognition in which the same picture of an apple, chair, or face is shown repeatedly up to 180 times for 250 ms each with an interstimulus interval of 100 ms in random order (see Supplementary Fig. S1a and Supplementary Table S1 for the number of trials of each subject after quality control). Each visual object is always presented together with a beep tone of the same duration, a low tone together with the apple, a middle tone with the chair, and a high tone with the face. The rationale behind this fast event-related approach was to maximize efficiency by delivering as many stimuli as possible within about three minutes while keeping stimulus durations long enough to preserve decodability. Catch trials with cartoon images and animations are interspersed after every first to fourth block of 36 repetitions to give breaks, keep up attention and maintain motivation (Fig. 1a). The subsequent second part serves to induce a sequential representation of the objects in memory. To this end, each of the three pictures is shown together with the corresponding beep tone in a fixed order, interspersed with catch trials. These triple-object sequences with a duration of 950 ms were repeated up to 50 times using a variable interstimulus interval of 300 to 600 ms (Fig. 1b). This way, time constraints were balanced against a sequence learning experience of sufficient intensity to create a durable memory representation strong enough to be captured with EEG. In the final part, which is embedded into the sequential part, infants did not rest passively but heard the low tone that was played in the previously learned sequence. This auditory stimulus, provided in large variable interstimulus intervals of 3,000 to 5,000 ms served as a cue to elicit and enhance forward reactivation of the object sequence (Fig. 1c). The length of resting-state recordings for each subject after quality control can be found in Supplementary Fig. S1b.

**Fig. 1.**
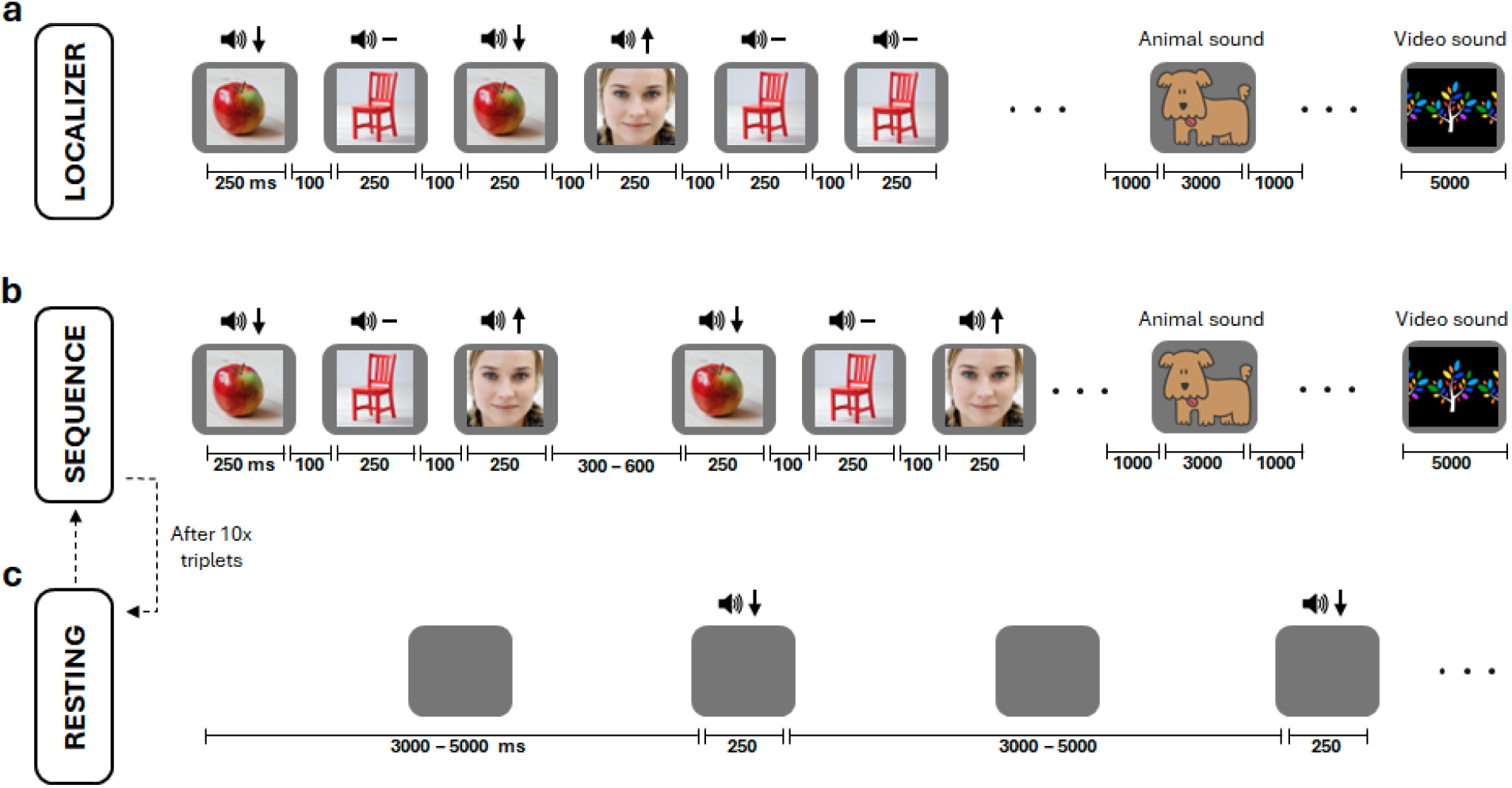
Experimental stimulation design. **a**, The localizer part comprises three different photo images shown together with a tone. Combinations were fixed: apple – low tone (down arrow), chair – mid tone (dash), and face – high tone (up arrow). Each of these images is presented twelve times per block. After each block, there is a short break during which a cartoon image of an animal together with a corresponding sound is presented. At every third break, a colorful video animation and an entertaining sound is presented instead. Blocks are repeated 18 times for a total of 180 repetitions per image. **b**, In the sequence learning part, the three images are shown in fixed order. These sequences are repeated eight times per block. Breaks are interspersed between blocks like in the localizer part. Blocks are repeated ten times for a total of 50 sequence repetitions. **c**, The resting-state part is embedded in the sequence learning part (dashed arrow), occurring after 10 repetitions of the triplet forming the sequence. During resting state, the screen remains empty and the low tone is played at variable interstimulus intervals to elicit and enhance the reactivation of the sequence of objects. These low tone cues are presented up to 50 times in total. ms = milliseconds. Images are license-free (sources: https://things-initiative.org and https://www.freepik.com).

### Time-resolved decoding of sequential reactivation

To determine the sequential reactivation of the objects presented, we first decoded short periods of infant’s resting-state data with a one-vs-rest logistic regression classifier trained on localizer data (Fig. 2ab). The resulting time courses of stimulus reactivation probability were then subjected to a temporally-delayed linear model^9,12^. This model allowed us to quantify the pairwise transition strengths from one stimulus reactivation to the other (e.g., from apple to chair) over a given time lag. Comparing matrices of these transition strengths to forward and backward sequence matrices (from apple to chair to face and vice versa), we obtained a measure of forward or backward sequential reactivation over time lags (Fig. 2c). A nuisance regressor was added to the matrices to rule out the possibility that the results were driven by oscillatory activity within the signal (see Methods for details).

**Fig. 2.**
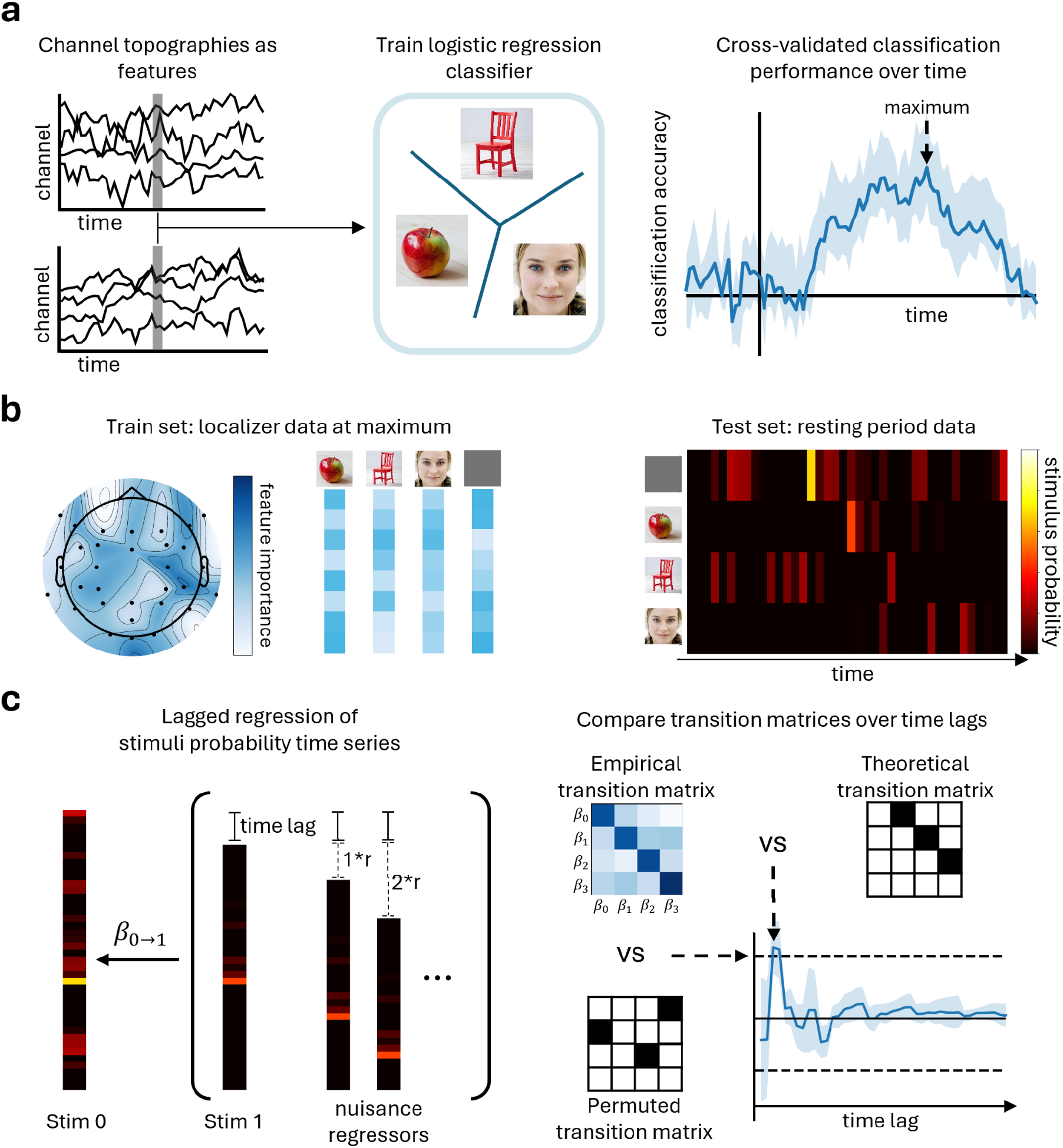
Decoding sequential reactivation at rest. **a**, The event-related EEG signal obtained from the localizer was compared between stimuli per time point by training a logistic regression model using six-fold cross-validation. Test set classification accuracies were averaged over folds and then over infants resulting in a grand average accuracy time series. **b**, To decode the resting-state data, another logistic regression model based on lasso regularization was trained on the time point of maximum classification accuracy in the localizer part (at around 400 ms). Training data included channel activations of all stimuli at the maximum time point and an additional baseline activity composed of channel activations at 100 ms before the actual stimulus onset (see Methods for details). The model was then tested on the infant’s resting-state data, transforming the raw activation into a time series of stimulus probability. **c**, Temporally-delayed linear modeling was then applied to the decoded resting-state signal, estimating one empirical transition matrix per time lag (in total: 50 evenly spaced time lags between 10 and 500 ms). Additional time-shifted copies of the stimulus probability time course (indexed by r) are added to the design matrix as nuisance regressors to control for oscillatory activity (see Methods for details). Each transition matrix was finally compared to a theoretical transition matrix (and their shuffled versions) resulting in a measure of sequenceness (and a permutation distribution of this measure) over time lags. The shaded areas displayed around the classification and sequenceness indices represent the standard error of the mean computed over participants (for details about temporally-delayed linear modeling, see Methods section *Decoding sequential reactivation*). All plots are based on simulated data to illustrate the procedures. ß = beta coefficient. Stim = stimulus. vs = versus.

### Group-level and individual reactivation sequences

Class labels of the three objects presented during the localizer experiment were decoded for each time point separately from trial-wise whole-brain EEG data using multivariate time-resolved decoding. This approach made it possible to create a feature space that captures the electrophysiological information carried by all recording electrodes instead of modeling each as a separate signal source. Classification was conducted individually for each participant and each object against the other two objects. Across participants, stimuli were classified significantly above chance level between 160 and 530 ms (Fig. 3a, p < 0.001, corrected, cluster-mass sign-flipped permutation test), reaching a maximum classification of around 60 % at 400 ms after stimulus onset. Individual time courses of each participant are shown in Supplementary Fig. S2. Spatial response patterns across participants were estimated for different time points. Stimuli were separable in bilateral occipital and posterior temporal electrodes (Fig. 3b).

**Fig. 3.**
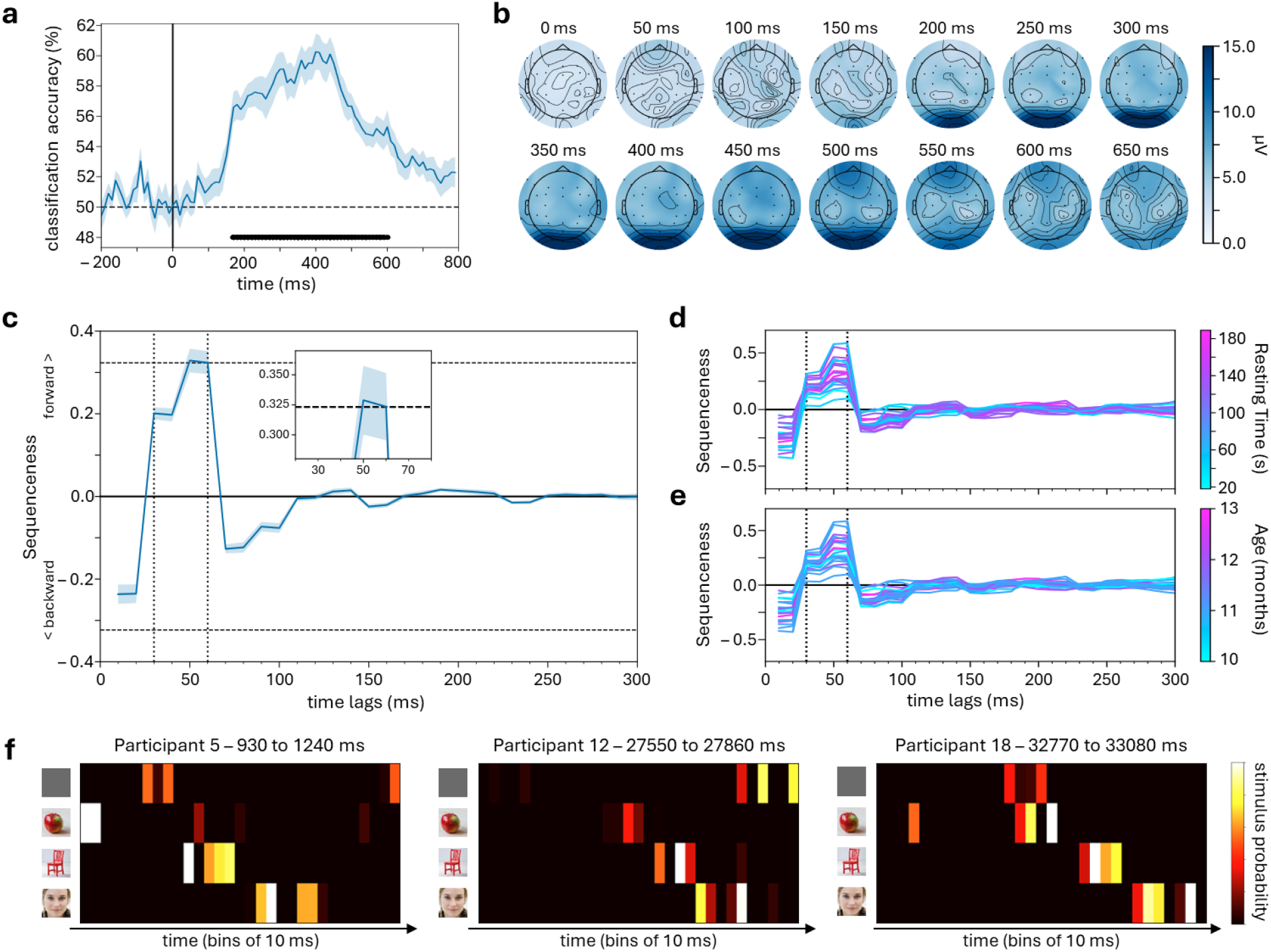
Group-level and individual reactivation sequences. **a**,**b** Localizer experiment. **a**, Group-level classification accuracy time courses. Cross-validated classification accuracy (y-axis), averaged across participants, is illustrated for each time point (x-axis). The shaded area depicts the standard error of the mean. The solid vertical line depicts the stimulus onset. The dashed horizontal line depicts chance level. Dots illustrate significant temporal clusters based on a cluster-mass permutation test (*p* < 0.001, sign-flipped). **b**, Electrode feature importance maps showing root mean square errors across class labels with higher values indicating larger deviation from the average EEG activity. Spatial electrophysiological response patterns were averaged across participants and calculated for several post-stimulus time points. Strongest feature importance for object discrimination was detected in bilateral occipital electrodes. The color bar denotes the standard deviation in microvolts (µV). **c–f**, Resting-state phase. **c**, Group-level sequenceness over time lags. A sequenceness index (y-axis) is plotted as a function of the average time lag (x-axis) between stimuli. Positive values indicate a forward sequence and negative values indicate a backward sequence. The curve represents the average across participants. Shaded areas depict the standard error of the mean. Dashed horizontal lines depict the maximum of all shuffled transitions across all time lags, corresponding to a corrected two-sided permutation threshold of p = 0.05. Dashed vertical lines serve as a visual aid highlighting time lags between 30 and 60 milliseconds. **d**,**e** Individual sequential reactivation over time. Axes correspond to **c**. The color bar in **d** denotes the time (s) each participant spent resting in the EEG. The color bar in **e** denotes the age (in months) of each participant. **f**, Examples of replay activity during the resting-state phase in three participants. Each bin along the x-axis corresponds to 10 milliseconds. Heatmaps depict normalized stimulus probability time series. ms = milliseconds.

At resting state, we detected a forward sequential reactivation effect at average time lags of about 50 to 60 ms between stimuli in the sequence (p < 0.05, two-sided, corrected, permutation test; Fig. 3c). This effect was consistent across individual participant data (Fig. 3d–f) (see Supplementary Fig. S3 for all empirical transition matrices for all time lags between 10 and 500 ms; see Supplementary Fig. S4 for separate beta values). There was no evidence for a significant association between sequenceness and the length of the resting time (*ρ* = 0.378, *p* = 0.101, Spearman’s rank correlation) or the age of the participants (*ρ* = 0.180, *p* = 0.447, Spearman’s rank correlation). There was also no correlation between the individual participant’s decoding performance in the localizer (at 400 ms) and the sequenceness (*ρ* = 0.008, *p* = 0.975, Spearman’s rank correlation). Moreover, cued replay events were distributed equally in time across participants (*D*(1946) = 0.02, p = 0.287; Kolmogorov-Smirnov test; Supplementary Fig. S5). It also turned out that the sequential memory representation is already formed during sequence learning at a timescale similar to the later cued reactivation phase (Supplementary Fig. S6).

As an additional control analysis, we applied the same approach for identifying sequential reactivation to two time periods occurring before the onset of sequence presentation. One time period included the break before the start of the localizer and the other time period included the break after the localizer. Both time periods did not reveal any significant effects of sequenceness (break before localizer: z = 1.58, p = 0.057; break after localizer: z = 0.87, p = 0.193; cluster-mass sign-flipped permutation tests; Supplementary Fig. S7). Additionally, at both times, the peak effect of sequenceness between 50 and 60 ms was significantly weaker than the corresponding effect of sequenceness after learning (after learning vs. before localizer: W = 20, p = 0.003; after learning vs. breaks within localizer: W = 13, p < 0.001; Wilcoxon signed-rank tests).

## Discussion

Here we developed a fast event-related stimulation paradigm allowing us to elicit the replay of a previously learned sequence of objects in infants from the age of 10 months while recording EEG. Multivariate decoding techniques were leveraged to capture the temporal information carried by the electrophysiological time courses. Capitalizing on this experimental framework, we discovered that awake resting human infants exhibit time-compressed and structured forward reactivation of a sequence of previously observed objects. Replay compressed the learned sequence in time by a factor of around 6 to 7. These results were consistent across individual participant data.

It is remarkable that temporal compression in infants was already within the range observed in adult rats in which compression factors of 5 to 20 have been reported^1–3^. A plausible prediction emerging from this finding is that fast replay in early life could enhance learning by increasing sequential memory capacity which might contribute to the superior sequential cognitive abilities of humans compared to rodents^22–24^. However, beside a number of differences between recording methods and analytical approaches, this result might also point to potential differences between spatial and non-spatial tasks that are beyond the scope of the present study. In non-spatial tasks, relatively high levels of compression, exceeding factor 60, have been found in human adults^12^.

Sequential reactivation patterns in infants could in principle originate from occipital recording electrodes. This location would be consistent with high-spatial-resolution data obtained from adult rats and humans^7,9,12,25^. However, the question whether replay had occipital cortical sources cannot be reliably answered by the current data set given the limited spatial resolution of the EEG. Similarly, an involvement of the hippocampal formation, as one would predict from multi-unit recording results in rats^7^, cannot be determined here. This is due to the fact that the sensitivity of electroencephalographic scalp sensors decreases exponentially with distance from the source of neural activity, which impedes the detection of deep hippocampal signals^26^.

The discovery that non-spatial replay is already operational in the infant brain suggests that such a fast reactivation mechanism would be a selective advantage for sequential learning in early life. Sequential learning is essential for the emergence of important cognitive functions, for example, in the domains of language and action^27,28^. A testable prediction that could guide this new research agenda is that the speed and stability of replay responses is positively related to sequential rule learning abilities which in turn might positively relate to language processing and action planning abilities. The first step, however, would be to confirm that cued replay also occurs in the auditory sensory modality given that infant brains that have access to memory contents in multiple sensory domains should have a particular selective advantage for sequential learning in inherently multisensory real-world contexts.

Here we collected a comparably large number of trials within a relatively short recording session of about 15 minutes in total. Following this strategy, we delivered a proof of principle demonstrating the feasibility of detecting cortical replay in individual infants. Accordingly, while the main contribution of this work is an advance in our current understanding of typical memory development, our study thus also holds promise to facilitate the translation of the approach into clinical settings. For example, the current results could stimulate further research on the early detection of neurodevelopmental disorders associated with learning and memory impairments^29,30^. A possible future direction emerges from the open question of whether infants at risk for autism exhibit altered sequential reactivation of social-emotional or language information (e.g., facial cues or syntactic structure). Furthermore, while the present findings suggest that cued replay is already present in infancy, longitudinal follow-up work is needed to directly trace the developmental trajectories of sequential memory capacities.

## Methods

### Participants

30 infants without conspicuous findings in medical early detection screenings were invited for one recording session. An age range of 10–12 months was chosen to be well ahead of the developmental stage at which infants start producing sequential combinatorial speech. While some children may enter this stage as early as around 14–15 months, multi-word utterances are typically not observed before 18 months^31,32^. At the same time, our objective was to ensure that participants were old enough to tolerate an experiment lasting about 15 minutes. Datasets of 10 participants were excluded because of either too much noise and movement artifacts in their data or because they did not generate any resting-state data (see below). A minimum target sample size of 20 was determined a priori based on previous publications in adults^9,12^.

Recording sessions were videotaped and valid trials in which the infant looked at the screen during stimulus presentation were determined by visual inspection of the videos by two raters. Interrater agreement across participants was high (Cohen’s κ = 0.783). Only trials identified as attended by both raters were included in the analysis. The remaining 20 infants (5 female, 15 male) had a mean age of around 11.3 months (range: 10–13 months; Supplementary Table S1a). All caregivers gave written informed consent to participate. The study was approved by the Ethics Committee of the Medical Faculty of the University of Leipzig, Germany (402/23-ek). Data acquisition took place at the Max Planck Institute for Human Cognitive and Brain Sciences in Leipzig, Germany, between September 2024 and April 2025.

### Stimulus material and experimental design

Three photo images, one depicting an apple, one depicting a chair, and another one depicting a woman’s face were selected for this study from the THINGS database (https://things-initiative.org/). These visual stimuli were shown against a gray background and located centrally on the screen, occupying a visual angle of approximately 16°. Images were always presented together with a beep tone of the same duration, a low tone together with the apple, a middle tone with the chair, and a high tone with the face.

The experiment consisted of three parts: a functional localizer, subsequent sequence learning and short resting-state phases. To maximize the number of trials, a fast event-related design was employed for both the functional localizer and the sequence learning. During the functional localizer, images were presented in blocks consisting of 36 trials with 12 repetitions per image. In each trial, the image was presented for 250 ms with an interstimulus interval of 100 ms before the next trial started, resulting in a presentation speed of around 3 Hz. To catch the infant’s attention, after every first to fourth block, cartoon images of animals (e.g., a dog) paired with an animal sound (e.g., bark) were presented for 3000 ms with a 1000 ms break before and after stimulus presentation. After every fifth block, a longer break was taken and accompanied by presenting an engaging video clip showing animals, trees, rainbows, etc. The whole localizer task comprised 15 blocks, so that each image was presented 180 times in a pseudorandom order, allowing for not more than 3 image repetitions in a row. During sequence learning, all three image-sound pairs were rapidly presented as triplets in a fixed order (i.e., apple, chair, face). Each image and sound appeared for 250 ms with a 100 ms interstimulus interval. Triplets were separated by short breaks that varied randomly between 300 and 600 ms. After every fifth triplet, a cartoon animal and a sound was presented, like in the functional localizer. After every tenth triplet, a short resting-state phase was recorded during which the screen remained blank for 16 to 26 seconds to measure spontaneous EEG activity. As a potential cue for eliciting replay memory activity during this time, the lowest tone of the sequence was presented randomly every 3000 to 5000 ms. A total of 5 tones was presented per resting block. After the resting block ended, a new sequence learning block started. In total, 10 sequence-learning and resting-phase blocks were presented. Additional breaks were introduced by the experimenter whenever the infant’s attention started to decrease. To ensure consistency across participants, decreasing attention was defined a priori as no fixation of the screen for about three seconds, a period corresponding to approximately three stimulus events during the localizer or one triple-object sequence during sequence learning. Recording was terminated on finishing the experiment or when the infant started to express discomfort. In case the experiment was terminated early, data was only included in the further analysis if the localizer and at least 2 sequence-learning and resting-state blocks were completed.

### Data acquisition

Infants stood or sat on their caregiver’s lap in front of the screen. Only data recorded while the infant was calm and not fussy were used for further analysis. All participants were continuously monitored by a camera. Room light was dimmed and room temperature was controlled to avoid sweating under the cap. Active electrodes (Brain Products GmbH, Gilching, Germany) were positioned using a custom montage including 34 electrodes (31 for electroencephalography: Fz, F3, F7, F9, FC5, POz, C3, T7, TP9, CP5, P7, P3, Pz, O1, O2, P8, P4, CP6, TP10, T8, C4, Oz, FC6, F10, F8, F4, Fp2, FC4, CP4, FC3, CP3; one for electrooculography; reference: FCz; ground: Fp1). The electrooculogram electrode was positioned on the right cheek. The electroencephalogram was recorded using a BrainVision Recorder (v. 1.22.0001, Brain Products) at a sampling rate of 1000 Hz with a low-pass filter of 250 Hz. Caps (Easycap GmbH, Wörthsee, Germany) were gelled before being placed on the head to minimize preparation time. Additional gel (with custom ingredients: hydroxyethyl cellulose, propylene glycol, sodium chloride, water) was applied to the electrodes until impedance was below a threshold of 50 kΩ. This procedure was limited in time to two to three minutes, depending on the compliance of each infant.

## Data preprocessing

Electrophysiological data were preprocessed using MNE (v. 1.8.1) (https://mne.tools/stable/index.html). Eye movements were monitored by two postdoctoral researchers with substantial EEG data collection experience (co-author C.M.P. and J.F.). Data inspection revealed that large eye movements and blinks most likely co-occurred with stronger effects of head and body movements during inattentive periods. These periods were removed from the analysis right away, and no further eye movement correction was applied on the remaining data since automatic signal correction using independent component analysis did not consistently identify eye movement-related components. Raw data were re-referenced to average activity before epoching.

For high-pass filtering, we used a trial-masked robust detrending procedure accounting for the fact that high-pass filtering can have an impact on the classification performance^33^. For the localizer, epochs were formed between 5 seconds before and 8 seconds after image onset. These large time windows are needed to reliably estimate low frequency trending activity in the data. The actual trial activity of interest between -0.2 to 0.8 seconds around the image was masked out (i.e., not included in the estimation) and trends were then iteratively estimated. Specifically, we first fit and removed a linear order 1 trend and then fit and removed a trend of order 30. Afterwards, epochs were cropped to the window of interest (-0.2 to 0.25 seconds around image onset). For the resting-state data, the robust detrending procedure was applied to the whole signal with a window size of 1 second. As before, first a linear and then a complex trend of order 30 were fit and removed from the signal. The detrended signal was then split into epochs of 3 seconds.

Individual trials in the localizer and resting-state data were automatically screened for artifacts using the *autoreject* package in MNE (https://github.com/autoreject). Optimal hyperparameter values determining the amount of channel-wise interpolation versus trial-wise rejection of signals were estimated for each infant based on a cross-validated grid-search over the parameter space. To this end, we specified four values for *n_interpolate* (1, 4, 8, and 16) and eleven evenly-spaced values for *consensus_perc* ranging from 0 to 1.0. To make sure that all artifacts were detected completely, both localizer and resting-state epochs were meticulously visually inspected afterwards. For the localizer, on average 51.59 % of trials (standard deviation: 16.94 %) were rejected per infant. For the resting-state, an average of 95.83 s of data (standard deviation: 42.73 s) were obtained. For further analysis, the signal was low-pass filtered below 50 Hz with an FIR filter and then downsampled to 100 Hz.

### Decoding the localizer data

For time-resolved decoding, voltage values were standardized for each channel and time point across trials. We then fit a logistic regression model with an L1 penalty on each time point separately. For the training and testing of the classifier, we used a 6-fold cross-validation procedure testing each stimulus against all other stimuli (one-vs-rest). The classifier underwent a balanced training procedure with stratified categories. Test accuracies per time point were then averaged across folds resulting in a time series of classification accuracy values. For each model, we also extracted the classifier weights and transformed them into spatial activation patterns to get channel-wise maps of feature importance for the classification^34^.

To test for significant above-chance classification performance, we used a cluster-based permutation test^35^. Specifically, we created a permutation distribution of group-average accuracies by randomly sign-flipping each infant’s accuracy time course around 0.5 and taking the average afterwards for 1000 permutations. In each permutation, clusters of consecutive significant activity over time were identified at a cluster-forming threshold of p < 0.001 based on a t-test against chance (chance-level = 0.5) while keeping only the largest cluster t-sum value. Comparing the cluster sizes of our actual data against this extreme cluster distribution effectively controls for an inflation of alpha error over time^36^. Clusters with t-sum values larger than 0.1 percent of the extreme data were deemed significant.

### Decoding sequential reactivation

A temporally-delayed linear model was employed to detect sequential stimulus reactivation during resting state. To this end, we created a custom-made Python implementation of the original Matlab code published by Liu and colleagues^9^.

To capture reactivation during resting state, we first trained a classifier on the localizer data (see section above). Specifically, from the average classification accuracy across infants (obtained using time-resolved decoding of the localizer) we selected the channel activations of all trials at the time point of maximum accuracy for our training data. In accordance with previous studies, we also added baseline activity taken from -0.1 s before stimulus onset to the training data^9,37^. The purpose of this step is to decorrelate classifier patterns and at the same time allow all stimuli to have low probability values. We included this baseline activity as a fourth stimulus in our model to increase the amount of possible permutations of the theoretical transition matrix (see below). For each infant, a logistic regression with an L1 penalty and a regularization strength of 1/6, following the literature standard^9,12^, was fitted to the standardized training data. This model was then applied to decode probability scores for stimulus reactivation (i.e., apple, chair, face, baseline) at each time point in the infant’s resting-state data.

For temporally-delayed linear modeling, stimulus probability time courses were entered into a lagged regression procedure in the following manner: For each combination of two stimuli, a regression model was fit with one stimulus probability time course as a dependent variable and a time-shifted copy of the stimulus time course as an independent variable. The time shift is denoted here as n where one n corresponds to a lag of 10 ms. To control for the influence of oscillatory activity in the signal, additional copies were added as nuisance regressors to the design matrix at n + r, n + 2r, n + 3r, etc., where r also corresponds to a lag of 10 ms and represents a control frequency (i.e., r = 10 would produce copies at n + 100 ms, n + 200 ms, n + 300 ms etc.). The estimated beta weight is therefore a measure of how reliably the reactivation of one stimulus follows the other at a specified time lag n. After running all comparisons between all four stimuli, this procedure yields a 4x4 empirical transition matrix of regression weights *β* per time lag where rows represent the starting position and columns represent the transition (where *β*_0→1_(*n*) is the transition strength from stimulus 0 to 1 at time lag n). Each empirical transition matrix was then entered as a dependent variable into a second regression with the following independent variables: (1) a binary matrix encoding the sequence of interest in the forward direction (i.e., 0→1, 1→2, 2→3; see Fig. 2c), (2) a transpose of this matrix representing the backward sequence, (3) the identity matrix to control for autocorrelation, and (4) an intercept. Subtracting the first two regression weights *z*_*forward*_ – *z*_*backward*_ then yields a measure that quantifies sequential reactivation in the forward direction (positive values) and backward direction (negative values). Here, we computed 50 evenly spaced time lags between 10 and 500 ms. As an oscillatory activity control, we chose r to be 8 (corresponding to a control frequency at n + 80 ms, n + 160 ms, n + 240 ms etc.). The rationale of controlling for oscillatory activity was not to account for the time scale of the trial as it includes multiple seconds of spontaneous resting signal but to control for alpha band activity. Specifically, if such rhythmic activity occurs in all signals, classical cross-correlation analyses would exhibit high correlations at a lag corresponding to the cycle length of the oscillation which could then be misinterpreted as sequential reactivation or potentially overshadow smaller sequential reactivation effects.

For statistical analysis, we generated a null distribution by permuting together both rows and columns of the theoretical transition matrix which resulted in 22 possible permutations (i.e., 4! = 24 matrices minus the matrices corresponding to the forward and backward sequence). For each permutation, we obtained one surrogate sequence measure over all lags per infant. All permutation data was averaged over participants and the average sequence measure was compared to the maximum value of the null distribution to obtain p-values.

The compression factor of replay activity is estimated as the fraction of the inter-stimulus interval during sequence learning (i.e., 350 ms) and a significant time lag identified by the sequenceness measure. For example, a significant time lag at 100 ms between stimuli would result in a temporal compression factor of around 3.5.

## Data availability

The data collected for this study will be made available after publication through a public link. During the peer review phase, data can be made available to the reviewers by the corresponding author upon request.

## Code availability

All code used for data analysis is available from GitHub (https://github.com/SkeideLab/REPLAY_code).

## Acknowledgements

This work was supported by the German Research Foundation (DFG Heisenberg Program Grant 433758790 awarded to M.A.S.) and the Jacobs Foundation (Research Fellowship awarded to M.A.S.). The funders had no role in study design, data collection and analysis, decision to publish or preparation of the manuscript.

## Author contributions

M.A.S. conceived the study. J.F. performed the experiments. C.M.P. analyzed the data and visualized the results with feedback from M.A.S. C.M.P. and M.A.S. wrote the manuscript. M.A.S. supervised the study.

## Ethics declarations

The authors declare no competing interests.

