## Supplementary Material for "Cued replay of memory contents in human infants"

### Supplementary Table S1

**Table S1.** Demographic information and the amount of included trials in the localizer and sequence presentation. Included trials are presented as total amount (N) and as a percentage of all trials in the data (% Retained).

| | | | Localizer | | Sequence | |
|---|---|---|---|---|---|---|
| Participant | Age | Gender | N | % Retained | N | % Retained |
| 1 | 12 | f | 146 | (27.0%) | 82 | (82%) |
| 2 | 11 | m | 248 | (45.9%) | 61 | (61%) |
| 3 | 12 | m | 443 | (82.0%) | 57 | (57%) |
| 4 | 11 | f | 352 | (65.2%) | 95 | (95%) |
| 5 | 13 | m | 266 | (49.3%) | 30 | (30%) |
| 6 | 11 | m | 223 | (41.3%) | 9 | (9%) |
| 7 | 12 | f | 277 | (51.3%) | 63 | (63%) |
| 8 | 11 | m | 196 | (36.3%) | 5 | (5%) |
| 9 | 10 | m | 257 | (47.6%) | 41 | (41%) |
| 10 | 11 | m | 348 | (64.4%) | 47 | (47%) |
| 11 | 10 | m | 352 | (65.2%) | 47 | (47%) |
| 12 | 12 | m | 91 | (16.9%) | 47 | (47%) |
| 13 | 13 | f | 349 | (64.6%) | 69 | (69%) |
| 14 | 11 | f | 177 | (32.8%) | 13 | (13%) |
| 15 | 12 | m | 300 | (55.6%) | 38 | (38%) |
| 16 | 11 | m | 208 | (38.5%) | 15 | (15%) |
| 17 | 10 | m | 305 | (56.5%) | 50 | (50%) |
| 18 | 11 | m | 224 | (41.5%) | 87 | (87%) |
| 19 | 11 | m | 492 | (91.1%) | 88 | (88%) |
| 20 | 11 | m | 348 | (64.4%) | 46 | (46%) |

### Supplementary Figures S1–S7

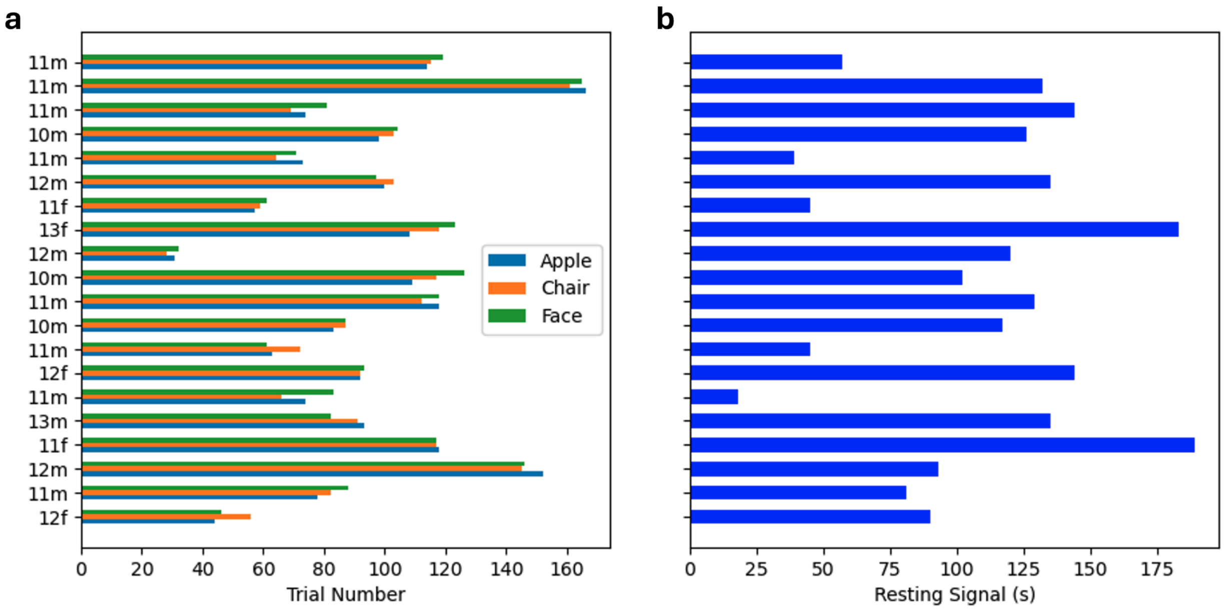


**Fig. S1 | Participant and dataset characteristics.** The x-axis of the left diagram depicts the number of trials in the localizer experiment after quality control. The x-axis of the right diagram depicts the length of the resting-state dataset. The y-axis refers to the age in months (10–13) and the sex (m = male, f = female).

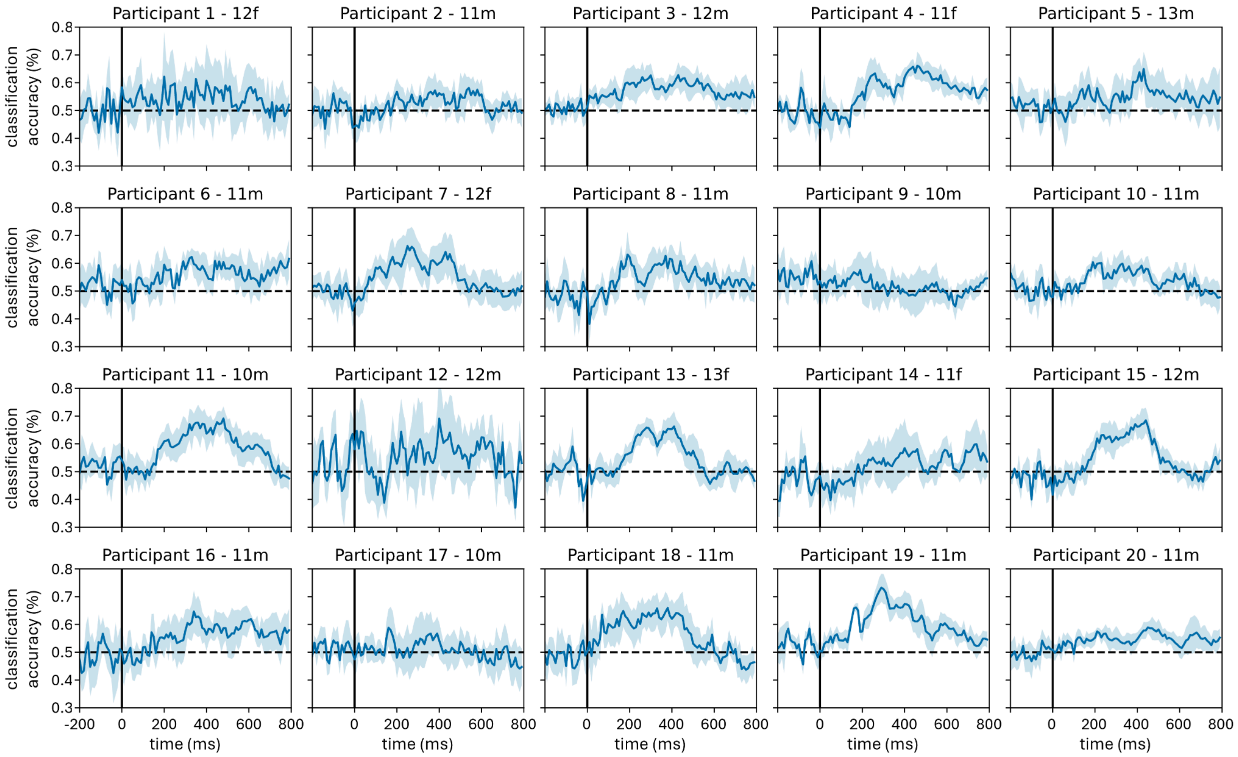


**Fig. S2 | Individual results of each participant obtained from time-resolved decoding of the localizer.** Cross-validated classification accuracy (y-axis) is illustrated for each time point (x-axis, ms = milliseconds). The shaded area depicts the standard error of the mean over six folds. The solid vertical line depicts the stimulus onset. The dashed horizontal line depicts chance level.

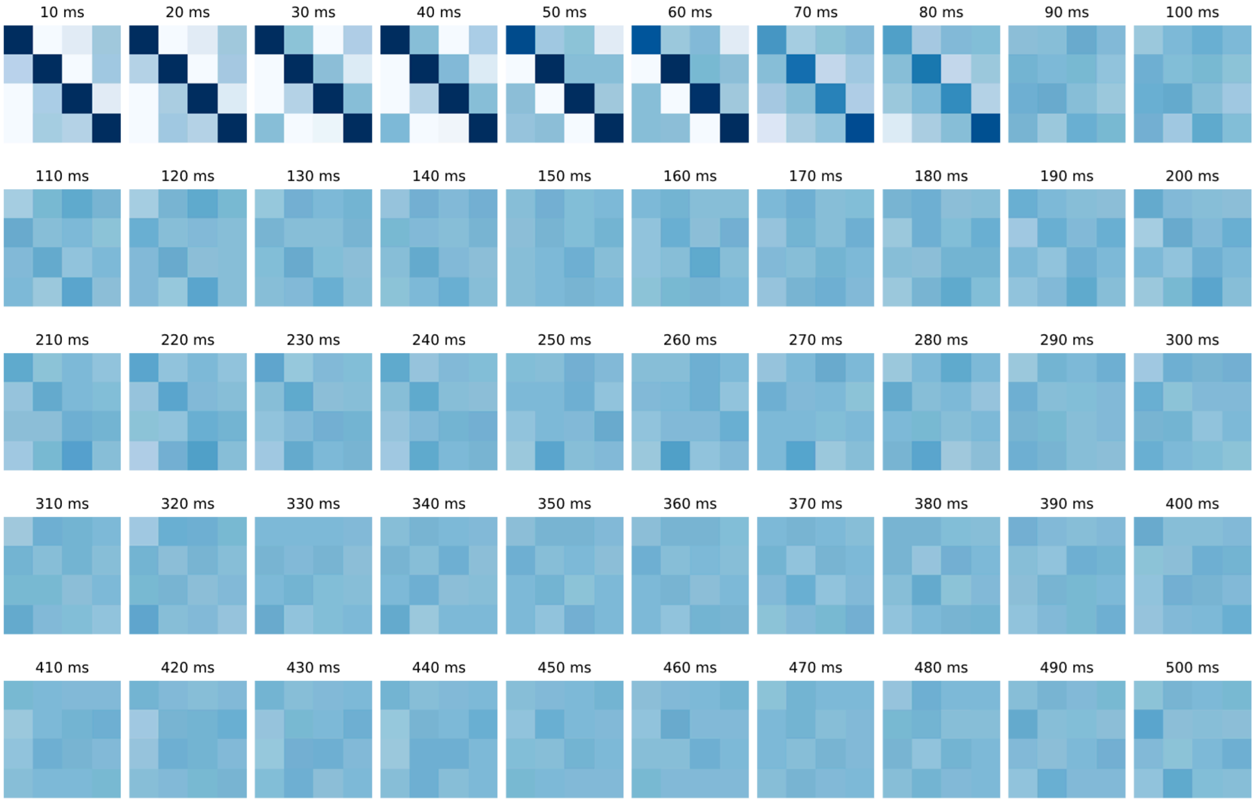


**Fig. S3 | Average empirical transition matrices.** These matrices show the beta values of the lagged regression on the stimulus probability time series. Within each lag, each beta value refers to a specific state transition encoded from columns to rows. The first row, for example, indicates the transition likelihood from baseline to each of the four states. On-diagonal values indicate the transition likelihood within each state. Matrices are averaged over subjects and color coded on the same scale.

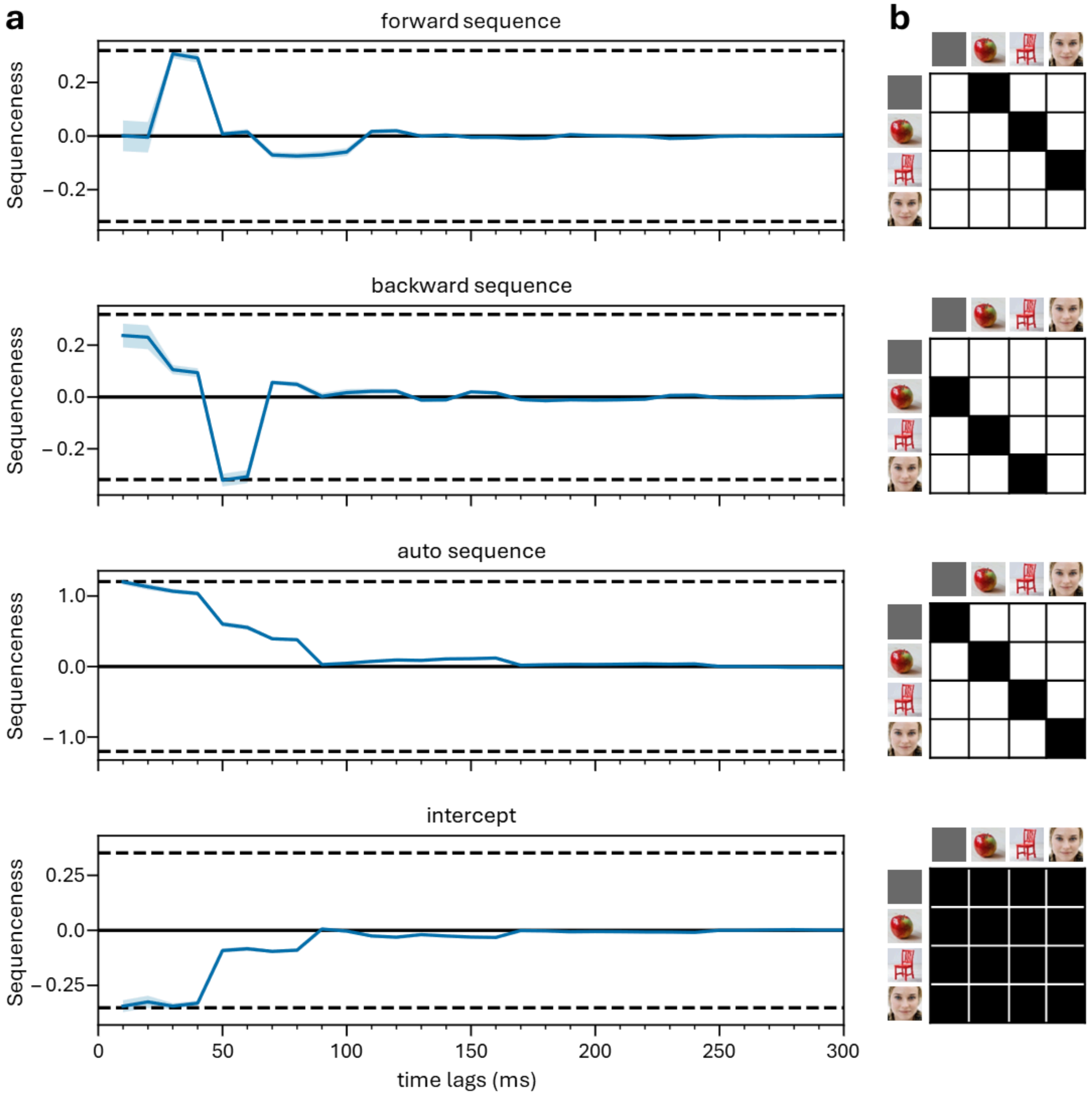


**Fig. S4 | Predictors of the empirical transition matrix. a**, The sequenceness index on the y-axis is plotted against the inter-stimulus time lag of the sequence (x-axis). Positive values indicate forward sequential reactivation and negative values indicate backward sequential reactivation. Shaded areas depict the standard error of the mean across participants. Dashed horizontal lines depict the 97.5 percentile of all shuffled transitions across all time lags, corresponding to a corrected two-sided permutation threshold of p = 0.05. **b**, Theoretical transition matrices used to fit the empirical transition matrix. Black cells indicate a value of one and white cells indicate a value of zero. State transitions are encoded from columns to rows (e.g., a one in the second column of the first row indicates a transition from baseline to apple). The auto sequence (i.e., the stability of the state over lags) and the intercept are control variables.

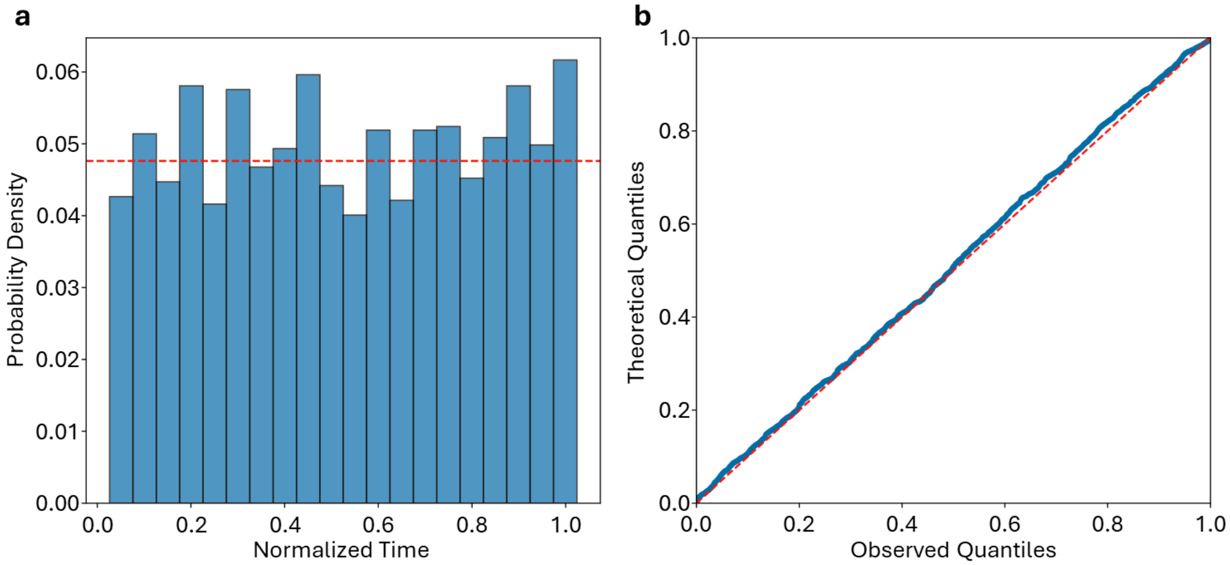


**Fig. S5 | Temporal distribution of cued replay events. a,** The histogram plot depicts the relative likelihood of cued replay events (probability density, y-axis) over time (x-axis). Time was normalized since the length of resting phases varied between participants. The dashed horizontal line marks the expected bin values of a uniform distribution. **b**, Quantile-quantile plot of cued replay events across all participants. The y-axis shows the quantiles of the theoretical distribution while the x-axis shows the quantiles of the sample data. The blue continuous line represents sorted replay events and the red dashed line indicates the expected uniform distribution.

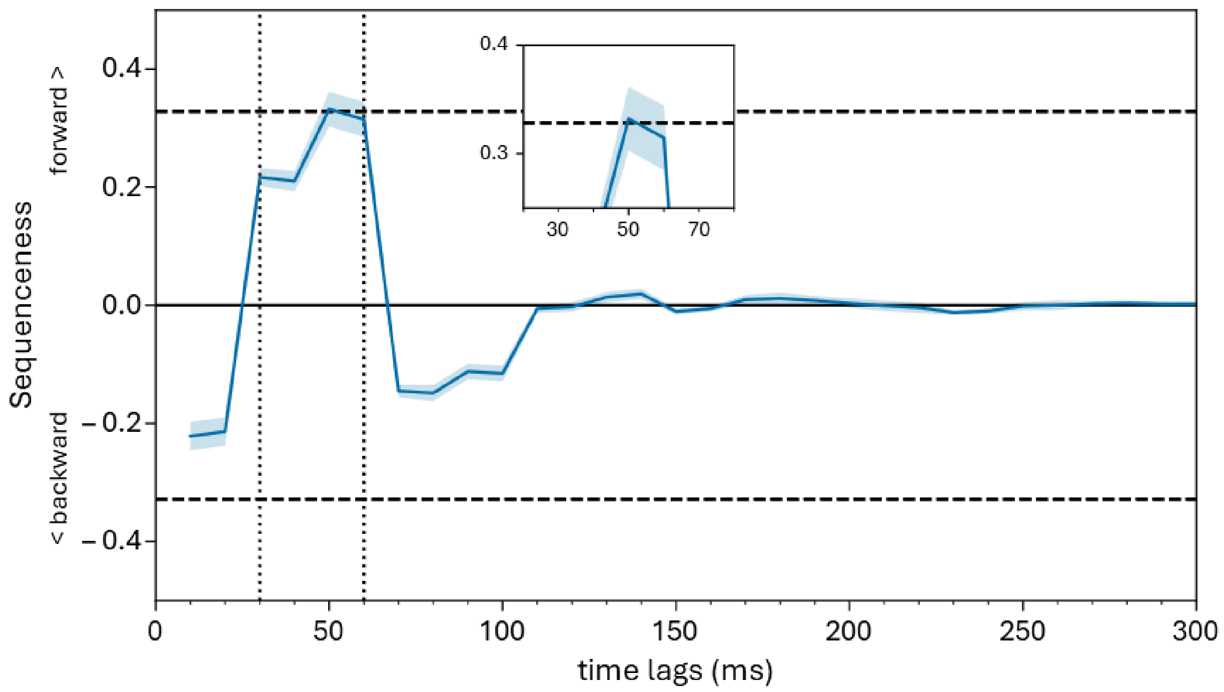


**Fig. S6 | Effect of sequenceness during sequence presentation.** The sequenceness index on the y-axis is plotted against the inter-stimulus time lag of the sequence (x-axis). Positive values indicate a forward sequence and negative values indicate a backward sequence. Shaded areas depict the standard error of the mean across participants. Dashed horizontal lines depict the maximum of all shuffled transitions across all time lags, corresponding to a corrected two-sided permutation threshold of p = 0.05. Dashed vertical lines serve as a visual aid highlighting time lags between 30 and 60 ms.

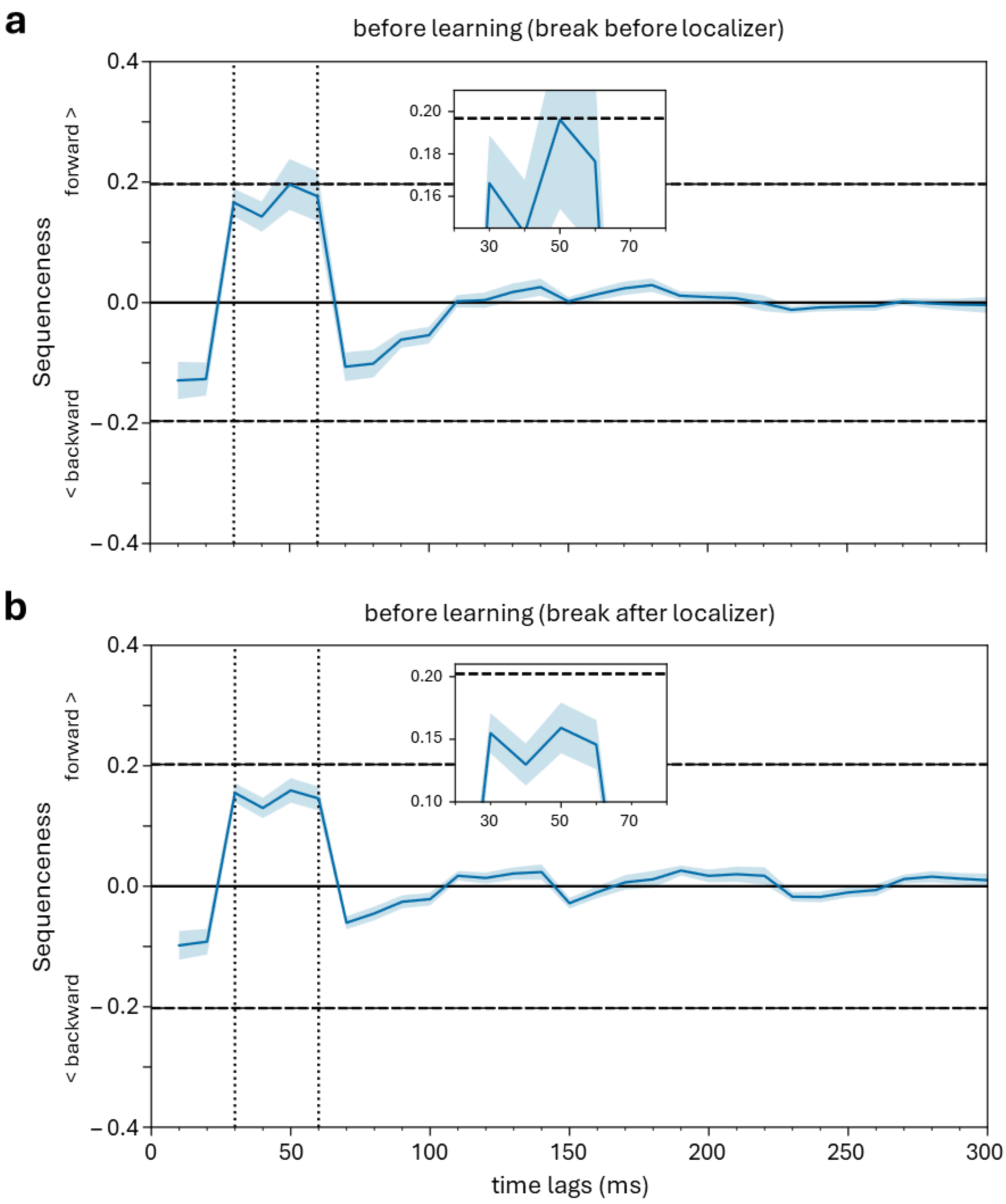


**Fig. S7 | No effect of sequenceness before sequence presentation. a**,**b,** In these control analyses, the sequence decoding approach was applied to two time periods preceding the onset of the sequence presentation: **a,** the break before the onset of the localizer, and **b**, the break after the localizer). The sequenceness index on the y-axis is plotted against the inter-stimulus time lag of the sequence (x-axis). Positive values indicate a forward sequence and negative values indicate a backward sequence. Shaded areas depict the standard error of the mean across participants. Dashed horizontal lines depict the maximum of all shuffled transitions across all time lags, corresponding to a corrected two-sided permutation threshold of p = 0.05. Dashed vertical lines serve as a visual aid highlighting time lags between 30 and 60 ms.
